# Have AI-Generated Texts from LLM Infiltrated the Realm of Scientific Writing? A Large-Scale Analysis of Preprint Platforms

**DOI:** 10.1101/2024.03.25.586710

**Authors:** Huzi Cheng, Bin Sheng, Aaron Lee, Varun Chaudary, Atanas G. Atanasov, Nan Liu, Yue Qiu, Tien Yin Wong, Yih-Chung Tham, Yingfeng Zheng

## Abstract

Since the release of ChatGPT in 2022, AI-generated texts have inevitably permeated various types of writing, sparking debates about the quality and quantity of content produced by such large language models (LLM). This study investigates a critical question: Have AI-generated texts from LLM infiltrated the realm of scientific writing, and if so, to what extent and in what setting? By analyzing a dataset comprised of preprint manuscripts uploaded to arXiv, bioRxiv, and medRxiv over the past two years, we confirmed and quantified the widespread influence of AI-generated texts in scientific publications using the latest LLM-text detection technique, the Binoculars LLM-detector. Further analyses with this tool reveal that: (1) the AI influence correlates with the trend of ChatGPT web searches; (2) it is widespread across many scientific domains but exhibits distinct impacts within them (highest: computer science, engineering sciences); (3) the influence varies with authors who have different language speaking backgrounds and geographic regions according to the location of their affiliations (Italy, China, etc.); (4) AI-generated texts are used in various content types in manuscripts (most significant: hypothesis formulation, conclusion summarization); (5) AI usage has a positive influence on paper’s impact, measured by its citation numbers. Based on these findings, suggestions about the advantages and regulation of AI-augmented scientific writing are discussed.

## 1 Introduction

The advent of AI in various sectors has marked a new era in the production and consumption of digital content (Brynjolfsson et al., 2023). Among the most notable developments is the rapid development of generative AI and large language models (LLMs). ChatGPT^1^, introduced in 2022, is an LLM based on GPT3 (Brown et al., 2020) with uncanny ability to generate text that closely mimics human writing. The ability of ChatGPT and similar models to produce coherent, contextually relevant texts has revolutionized content creation, leading to its adoption across multiple writing forms. This proliferation has not been without controversy, however, as it raises significant concerns regarding the authenticity, originality, and quality of AI-generated content (Brynjolfsson et al., 2023; Cardon et al., 2023; McKee and Porter, 2020; Salvagno et al., 2023). Moreover, the potential of these technologies to contribute to information overload by producing large volumes of content rapidly has been a subject of debate among academics and industry professionals alike (Jakesch et al., 2019; Dergaa et al., 2023).

In the scientific community, the penetration of AI-generated texts poses unique challenges and opportunities. Scientific writing is typically characterized by its rigorous standards for accuracy, clarity, and conciseness, and some of these tasks could be assisted with LLM and AI-generated text. However, scientific writing also requires the art of human inquisition, perception, and a nuanced understanding and explanation of the key observations and findings; these parts of the scientific writing are not currently possible with LLM models. Thus, scientific writing may be a crossroads with the integration of AI-generated content. The core of this study focuses on exploring the extent to which AI-generated texts have made their way into scientific literature, particularly within the domain of preprint manuscripts. By leveraging a large open composite dataset of preprint submission and advanced detection tools, such as the Binoculars LLM-detector (Hans et al., 2024), this research aims to map out the landscape of AI influence in scientific writing. Our investigation spans across different disciplines and examines the correlation between the surge in AI-generated content and various factors, including search trends, domain-specific impacts, and the demographic characteristics of authors. We also examined the relationship between a paper’s impact and its AI usage, revealing that AI usage positively correlated with citation numbers. This comprehensive analysis provides insights into how AI is reshaping the conventions of scientific writing and offers more fine-grained suggestions about safe use of AI in academic research.

## 2 Results

### 2.1. Dataset

The publishing life-cycle of a paper may take various time periods, with some of them longer than a year. On the other hand, LLM-based text generation AI tools like ChatGPT, have gained broad popularity since the end of 2022. Such a short time span makes it difficult to analyze the AI footprint in officially published literatures. Therefore, we have instead focused on manuscripts submitted to preprint platforms. Such platforms like arXiv are good choices for our purpose for several reasons: First, more and more authors tend to upload a preprint version before they submit the manuscript to a journal to plant a flag about the timing of their discovery. Thus, the timeliness of content may be the latest we can get from the science community; Second, the number of manuscripts submitted to these platforms are high even in a short interval, making it possible to do more fine-grained analysis; Last, to our knowledge, all preprint platforms are open to bulk access, making large scale analysis possible.

In this study, we collected manuscripts in the form of PDF files from three mainstream preprint platforms: arXiv, bioRxiv and medRxiv (Fig. 1A, Fig. 1C left), covering domains spanning from math and engineering to biology and medicine (Fig. 1C middle). For all platforms, we downloaded manuscripts from 2022.01.01 to 2024.03.01. We chose this time period because it includes one year before and one year after the release of ChatGPT in December 2022. For each month, at most 1000 random manuscripts (in some months medRxiv has fewer than 1000 papers submitted) were downloaded in each platform using the provided API. After cleaning and preprocessing (see Method), some invalid documents were removed and 45129 manuscripts were used for analysis. The domains of these papers are categorized into following classes: Biological Sciences, Computer Science, Economics and Finance, Engineering, Environmental Sciences, Mathematical Sciences, Medicine, Neurosciences, and Physical Sciences.

**Figure 1:**
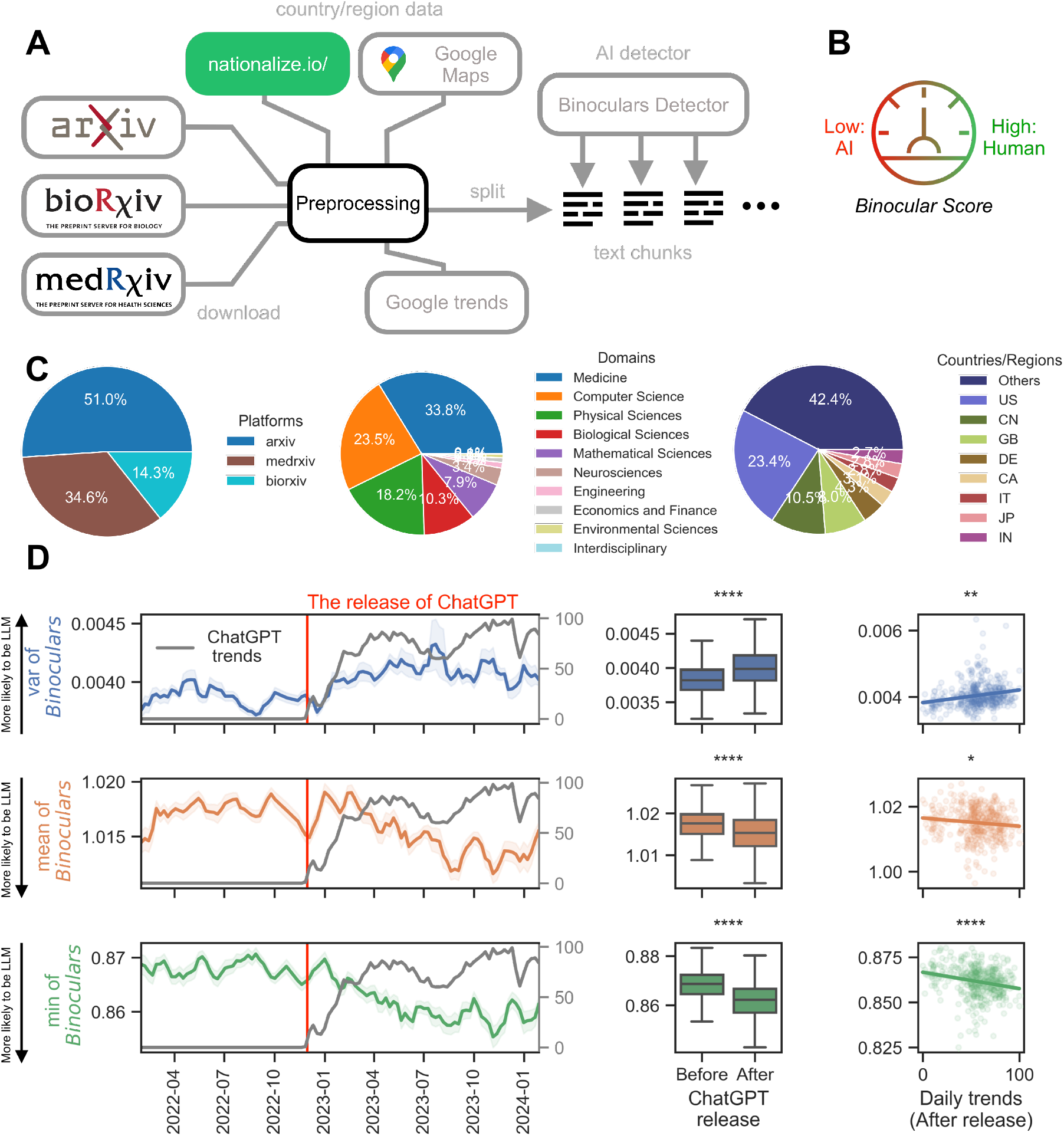
Overview of the data processing pipeline and analyses of AIs influence on scientific literature A: Schematic of the data processing workflow. Manuscripts and metadata were downloaded from three platforms: arXiv, bioRxiv, and medRxiv. Next, country/region information was extracted from the metadata using APIs provided by services such as nationalize.io and Google Maps. Following this preprocessing process, manuscripts were segmented into chunks. These chunks were then analyzed using Binoculars detectors to obtain chunk-level Binoculars scores. B: Interpretation of the Binoculars score. A higher Binoculars score suggests human authorship and a lower score indicates potential AI generation. C: Summary of dataset characteristics including the distribution of manuscripts across different preprint platforms (left), domain diversity within the dataset (middle), and country/region distribution (right). D: Examination of the AIs influence before and after the release of ChatGPT. Left: The weekly level comparisons between ChatGPT Google Trends and Binoculars indices. Top: the average variance of Binoculars values; Middle: the average of mean values of Binoculars; Bottom: the average of min Binoculars values. Middle: daily level comparisons of Binoculars indices before and after the release of ChatGPT. Right: daily level comparisons between ChatGPT Google Trends and Binoculars indices, after the release of ChatGPT. (For all statistical tests performed in D,* : *p* < 0.05, ** : *p* < 0.01, *** : *p* < 0.001, **** : *p* < 0.0001)

On the other side, since we have no internal access to the specific traffic data of OpenAIs website, to investigate the influence and usage of ChatGPT, we used Google Trends data as a proxy (Nuti et al., 2014). Daily and Weekly level world-wide Google Trends of the keyword ChatGPT are used for analyses at different temporal resolutions.

### 2.2. Binoculars scores before and after the release of ChatGPT

In early models like LSTM or GRU (Hochreiter and Schmidhuber, 1997; Cho et al., 2014), machine-generated texts could be easily spotted and were generally considered useless in production. However, since the advent of transformer-based models, detecting AI-generated texts has become challenging due to the transformative power of the architecture. The release of ChatGPT at the end of 2022 further complicated detection, as detectors may not have access to the model. On the other hand, LLMs like ChatGPT can generate seemingly realistic texts at first glance, making catch-by-eye detection implausible. Detectors that use hidden statistical patterns have become advantageous in this context, as they require no knowledge of the specific LLMs used and little to no training at all.

Some common choices are based on the perplexity of the given text (Dhaini et al., 2023; Ghosal et al., 2023; Tang et al., 2023). The general idea behind this approach is that texts generated by LLMs tend to have lower perplexities. However, this may only work for texts that are completely generated by LLMs. In the case of scientific writing, authors may rely on LLMs more for revising content rather than using LLMs to generate an entire manuscript from scratch. Detecting such revised texts could be extremely challenging. We noted that a tool developed recently, the Binoculars score (Hans et al., 2024), specifically addresses this issue. When Binoculars is high, it indicates that the input text is more likely generated by humans. When Binoculars is lower than a certain threshold, the text is more suspicious of containing LLM-generated content (Fig. 1B). By utilizing two instead of one LLM, Binoculars allows us to detect texts that may have prompts mixed into the content. This feature enables it to outperform several other known LLM detectors, such as Ghostbuster (Verma et al.,2023), GPTZero^2^, and DetectGPT (Mitchell et al., 2023), in many benchmark tests, including datasets involving arXiv samples. Given its outstanding performance efficiency, we use it as the main tool for detecting LLM-generated texts in this study (for details, see Material and Methods, Supplementary Materials).

Since a complete manuscript is usually too long for a single pass of Binoculars detector, we first split each manuscript into even-sized chunks. Then, each chunk is fed into the Binoculars detector, and a Binoculars score is calculated. For a manuscript, its LLM fingerprint is then the sequence of the corresponding Binoculars scores. In our study, we observed that the mean, variance, and minimum values of this sequence are crucial for spotting AI-generated texts. For all manuscripts in the dataset, we calculated their paper-level mean, variance, and minimum Binoculars scores. Next, a forward rolling average of these three scores with a window of 30 days was used to compute three Binoculars indices from 2022 to 2024, assuming the current usage of ChatGPT may be reflected in the manuscripts submitted in the near future rather than at the same moment, given that a manuscript usually takes a relatively long time to finish (a 30-day average lag is assumed).

We then compared these three indices with the weekly Google Trends of the keyword ChatGPT, which is used to indirectly measure the usage and popularity of AI tools in writing. As shown by the gray lines in the left column of Fig. 1, the search trend for ChatGPT rises after its release on 2022.11.30. Compared with this trend, we noticed that the three Binoculars indices correlate with the trend in various ways: The average mean and minimum Binoculars values are higher before the release of ChatGPT, while the variance is higher after the release. This suggests a divergence in content generated by humans and ChatGPT after the release, given the increase in variance and minimum value. The overall content containing ChatGPT-generated text is also higher, as indicated by the decrease in mean Binoculars indices.

We further examined whether this relationship holds in an even more refined temporal domain. Similarly, daily-level ChatGPT Google Trends were compared with Binoculars indices at the same resolution, but only for the time after the release of ChatGPT. The results on the right side of Fig. 1D indicate that the correlation persists and is consistent with the weekly level analysis. A closer look at the correlation significance reveals that, compared with mean Binoculars scores, minimum values and variances are more representative. This matches the impression we got from the weekly data that minimum values are stronger signs of the Google trend. It also implies the same aforementioned divergence tendency and the increased variance could be mainly driven by the increased min values. Therefore, in the later analysis we focus on the min and mean values of Binoculars indices.

### 2.3. Domains

The results in Fig.1D lay the ground for a more detailed analysis. We next ask: Is there a difference in the use of ChatGPT or other LLMs across different domains? If so, several factors could contribute to it: The distribution of corpora used for LLM training might be imbalanced, leading to differentiated performance across various domains, which in turn affects domain-specific usage preferences. For example, domains like mathematics, which often involve more abstract descriptions and highly contextualized symbols, may find it challenging to use ChatGPT directly, potentially resulting in higher Binoculars scores. The reliance on and familiarity with the latest digital tools may also lead to varying attitudes towards using LLMs in writing. For instance, the computer science community might be more open to integrating ChatGPT into their workflow.

To understand this, we categorized all manuscripts into several domains (Fig. 1C middle) and analyzed the distribution of mean and minimum Binoculars scores before and after the release of ChatGPT. Fig. 2A reveals the existence of the hypothesized differentiation. Domains such as biological science, computer science, and engineering show the largest drop in minimum Binoculars values after the release of ChatGPT, suggesting a relatively heavy use of it. In the domains of engineering and computer science, the mean Binoculars scores also significantly decrease. However, these domains also exhibit a relatively low average Binoculars score even before the release, which may be attributed to the size of the corpus of these domains in the training dataset of LLMs like ChatGPT. All other domains also demonstrate a decrease in either mean or minimum Binoculars scores, suggesting a widespread use of ChatGPT in scientific writing after its release.

**Figure 2:**
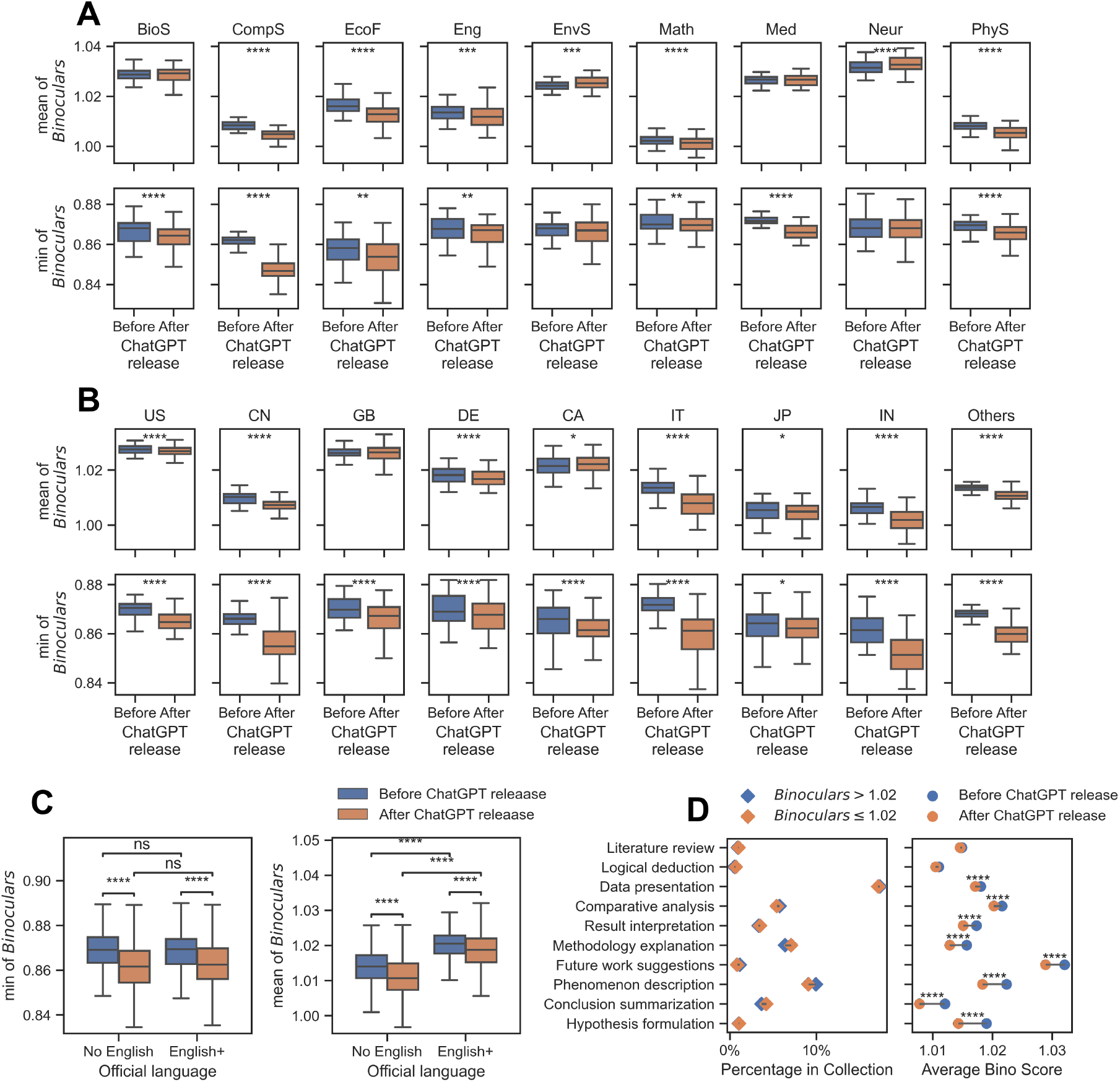
Distributions of Binoculars mean and min values across different domains (A), countries/regions(B), authors languages (C), and content types (D). A: Mean (top) and min (bottom) values across scientific domains before and after the release of ChatGPT. For each domain, the distribution of Binoculars scores is compared between manuscripts submitted before and after the release of ChatGPT on November 30, 2022. Domains are represented by their abbreviations: Biological Sciences (BioS), Computer Science (CompS), Economics and Finance (EcoF), Engineering (Eng), Environmental Sciences (EnvS), Mathematical Sciences (Math), Medicine (Med), Neurosciences (Neur), and Physical Sciences (PhyS). Asterisks denote the level of statistical significance for the difference in Binoculars scores before and after ChatGPT’s release within each domain. B: Same as A, but for different countries/regions. Note that besides the top 8 countries, the “other” column aggregates the results from all other countries/regions in the dataset. C: Similar to A and B, but for countries/regions with and without English as an official language before and after the release of ChatGPT. The box plots show the distribution of Binoculars scores for manuscripts from countries/regions where English is an official language versus those where English is not an official language. D: Analysis of content types in relation to AI-generated text. Left: Distribution of content types in texts with high (Binoculars scores above the average score of 1.02) and low (Binoculars scores below the average score of 1.02) Binoculars scores; Right: Distribution of content types before and after the release of ChatGPT, ordered by descending differences, from top to bottom. (For all statistical tests performed,* : *p* < 0.05, ** : *p* < 0.01, *** : *p* < 0.001, **** : *p* < 0.0001)

### 2.4. Countries/languages

Another important factor that may influence the use of ChatGPT is the native language spoken by the authors of the paper (El kah et al., 2023; Hwang et al., 2023). Since most of the manuscripts analyzed and published are in English, it is reasonable to hypothesize that individuals who use English as a second language may rely more on ChatGPT. However, directly analyzing this is impractical, as platforms do not provide the nationality of all authors. Additionally, an author may be fluent in more than one language. For each platform, we devised workarounds to address this issue (for details, see Methods) and therefore assigned a country/region for each manuscript in the dataset (Fig. 1C right). Top 8 countries with highest number of submissions are selected for analysis. For remaining countries/regions, we aggregated them into the “Others” category.

Similar to Fig. 2A, we analyzed the distribution of mean and minimum Binoculars values before and after the release of ChatGPT. From Fig. 2B, it is evident that almost all countries exhibit a decrease in minimum Binoculars values, while the decrease in mean Binoculars values is present but not as pronounced. Additionally, countries like China, Italy, and India show a larger gap in both the mean and minimum Binoculars values drop after the release of ChatGPT. We hypothesized that this is related to the fact that the native languages in some of these countries do not include English.

To validate this hypothesis, we classified countries/regions by their official languages (Fig. 2C). The results show that, although Binoculars scores decrease for all after the release of ChatGPT, the overall levels of mean and minimum Binoculars values are still higher in countries/regions where English is one of the official languages. This finding aligns with some previous studies indicating that some LLM detectors tend to recognize texts written by non-native English speakers as LLM-generated (Liang et al., 2023).

### 2.5. Content types

Another aspect to consider is the influence of AI-generated text on content types. Intuitively, content that introduces previous findings or contains more existing information might be more influenced by AI, as the training dataset could already have the knowledge. In contrast, highly specific content and new findings might be less suitable for generation by AIs. To examine this, we used the NLI-based Zero Shot Text Classification model to categorize all chunks in each manuscript into these types: phenomenon description, hypothesis formulation, methodology explanation, data presentation, logical deduction, result interpretation, literature review, comparative analysis, conclusion summarization, future work suggestions, bibliography, and publishing metadata. Excluding the last two types, which are irrelevant to our analysis and are introduced by the parsing process of PDF files, we analyzed the distribution of all these types in the dataset.

Specifically, we first checked if the distribution of content types is stable across text chunks with high and low Binoculars score distributions (Fig. 2D, left). All text chunks were split into two sets: those with Binoculars scores higher than the average score of the whole dataset (1.02) and those that were lower. We found that the average Binoculars scores of different content types matched our intuition: literature review content has very low Binoculars scores, while contents containing novel information, such as data presentation and phenomenon description, have the highest average Binoculars scores. Additionally, the content type distributions in high and low Binoculars score collections are relatively stable, as the portion shifts are small.

Next, we examined the Binoculars score differences for each content type before and after the release of ChatGPT (Fig. 2D, right). Although most content types showed signs of a decrease in Binoculars scores, literature reviews did not experience a significant drop after the release of ChatGPT. Contents previously considered “novel,” such as hypothesis formulation, conclusion summarization, phenomenon description, and future work suggestions, instead show the largest score drops.

### 2.6. Binoculars score and paper’s impact

One of the common reasons people worry about the use of AI is that it may “contaminate” content quality, but is this really the case? Since directly accessing such a subjective measure is hard, we turned to citation numbers as a proxy for a paper’s impact. Using the API provided by Semantic Scholar, we collected citation numbers for nearly all manuscripts in the dataset. We first compared the correlation between the Binoculars score mean values and the number of citations in two sets: manuscripts submitted before and after the release of ChatGPT. The correlation before the ChatGPT release is not significant (0.004214, p=0.56). However, after the release, the correlation changes to -0.018911, with a p-value of 0.002566. A difference in correlation analysis shows that the change in correlation is significant (p-value = 0.007994). This surprisingly implies that since people can use ChatGPT, the more one uses it (lower Binoculars score mean values), the more likely one will get citations.

To rule out the possibility that this difference is caused by the time elapsed effect on citations, we conducted a more fine-grained analysis. We first noticed that the distribution of citation numbers is highly imbalanced, with most papers receiving a few citations (Fig. 3A). Besides, citation numbers naturally accumulate, and thus more recent papers usually get fewer citations compared with older ones, though their actual impact may be comparable. This is reflected in the “decaying” daily citation average of citation numbers in the dataset (Fig. 3B). With such heterogeneous distributions, we compared correlations between the mean Binoculars scores and citations in a 30-day period, ranging from 2022 to 2024 (Fig. 3C left), as the accumulation effect could be ignored in such a short interval. The results show that, after the release of ChatGPT, there’s indeed a declining trend of this correlation down to negative regions. The difference in these month-level correlations is also significant, being consistent with our preliminary analysis above.

**Figure 3:**
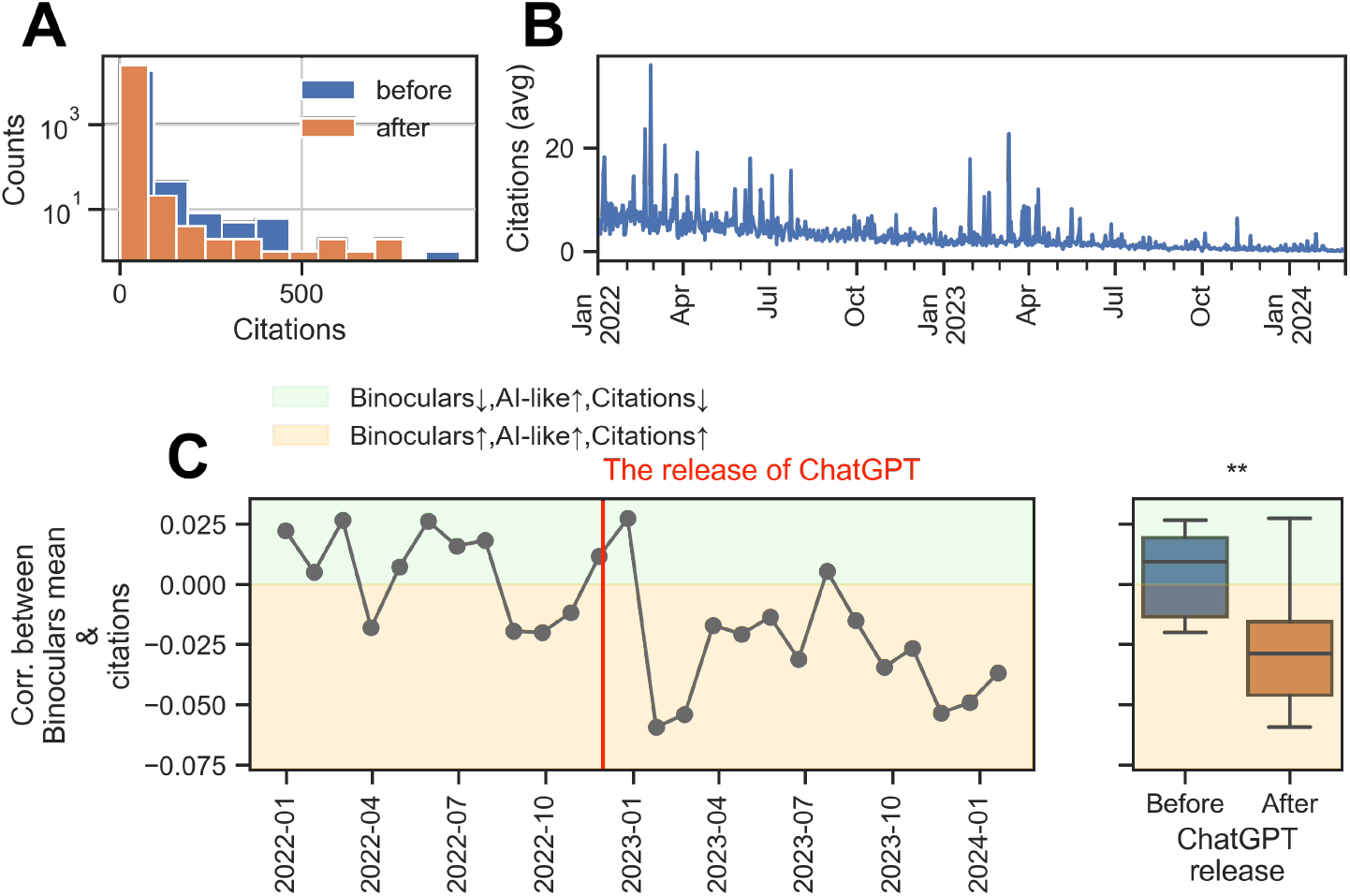
A: The histogram of manuscripts’ citation number before (blue) and after(orange) the release of ChatGPT. B: The daily average citation numbers in the whole dataset across all preprint platforms. C: The 30-day correlations between a manuscripts’ mean Binoculars score and their citation numbers from 2022 to 2024 (left) and the aggregated distributions of 30-day correlations before (blue) and after(orange) the release of ChatGPT (right). The light green regions indicate that when a manuscript more likely contains AI-generated text, it has a lower Binoculars score mean and less likely receives citations (positive correlation). For the light blue regions the trend is flipped due to the negative correlation.

## 3 Discussion

By analyzing around 45,000 manuscripts submitted to 3 preprint platforms over the past two years, we have identified a significant increase in the use of AI in scientific writing following the release of Chat-GPT in late 2022. This was achieved by examining the Binoculars score statistics for each manuscript. We observed that the average Binoculars scores have significantly decreased after 2022.11.30, and this decrease correlates with the Google Trends data for the keyword “ChatGPT “, indicating a widespread presence of AI-generated text in scientific manuscripts. Further analyses reveal an imbalance in AI usage across different disciplines and countries. Fields like computer science and engineering show a higher incidence of AI use. A similar trend is observed in countries where English is not the official language, as confirmed by integrated Ordinary Least Squares (OLS) regression analysis (see Material and Methods, Supplementary Materials). The influence of AI on content type is also uneven. Texts that are believed to contain new information exhibit a larger decline in Binoculars scores compared to literature reviews. Additionally, we have tracked the evolution of correlations between the mean Binoculars scores and citation numbers each month. An unexpected trend reversal was noted: before the release of ChatGPT, the correlation was weakly positive and at times insignificant, suggesting that writing style had a minimal impact on a paper’s influence. However, post-release, the correlation turned negative, indicating that papers with AI-generated content are more likely to be cited.

Nevertheless, the analysis pipeline constructed still has a few limitations. First, as pointed out by Hans et al. (2024), it is impossible to completely determine whether a text is generated by AI. Since the Binoculars score relies on statistical patterns more commonly found in AI-generated texts, theres a chance that improper use and attacks may reduce its reliability (Sadasivan et al., 2023). This is also why we observe fluctuations in Binoculars values in manuscripts before the release of ChatGPT. Other statistical tools we used for analysis, like the zero-shot text classification model, also make similar errors. Second, although an increasing number of authors tend to upload their manuscripts to preprint platforms (Piwowar et al., 2018), these platforms still do not cover all scientific papers, not to mention that different domains also have varying tendencies in using preprints as a distribution channel. Therefore, the dataset we used cannot represent the entire picture, though the statistical results are stable for major domains and countries/regions. Third, due to the limitations of platforms like arXiv, we have no direct access to the authors’ country/region/native language information. The introduction of nationality inference services inevitably leads to errors in specific papers. Moreover, as discussed above, a manuscript may contain contributions from people speaking different languages, making the country/region analysis imprecise.

Despite the limitations outlined above, this study represents, to our knowledge, the first endeavor to reveal the trend of AI’s footprint in contemporary scientific writing activities through quantitative, large-scale analysis. Regardless of personal opinions, AI tools such as ChatGPT have become embedded in daily human communication. However, unlike scenarios where students employ AI to complete assignments, scientific writing has traditionally been viewed as a means of disseminating ideas and new knowledge to humanity, raising concerns within the community about ethical issues more than just plagiarism (Sadasivan et al., 2023).

Our analysis suggests that, for the regulation purpose, the impact of AI on scientific writing should be discussed at more detailed levels, rather than simple usage disclosure. First, although the mean Binoculars scores have significantly decreased following the introduction of ChatGPT (Fig. 1D), they remain above 1.01, which is considerably higher than the threshold of around 0.9 set by (Hans et al., 2024). This suggests that while the use of AI in scientific writing may be widespread, it is not predominantly for generating extensive texts. Authors may primarily use AI for editing and revising purposes, especially in content that is about the creation of new knowledge (Fig. 2D). With such applications, we cant see any reasons to simply oppose the use of AI in writing, as it can be used to bridge the communication gap caused by region/language barriers. The heterogeneous use of AI in countries without English as their official languages indirectly confirmed this point (Fig. 2C). Analysis about the correlation between Binoculars scores and citation numbers, on the other side, also suggests the positive influence of AI usage in improving papers’ impact (Fig. 3C).

## 4 Material and Methods

### 4.1. Data source and preprocessing

To extract submitted manuscript information from bioRxiv and medRxiv, we used the official “details” API in these platforms https://api.biorxiv.org/ and https://api.medrxiv.org/. For arXiv, manuscript information is downloaded directly from Kaggle (https://www.kaggle.com/datasets/Cornell-University/arxiv) as it provides free bulk access to all submissions on arXiv. For all platforms, we collect at most 1000 submissions in each month, starting from 2022.01 to 2024.03.

All PDF files were directly download from the corresponding platforms using the URLs in the meta data fetched above. Once downloaded, we used pymupdf (https://pymupdf.readthedocs.io/en/latest/) to parse the PDF files and turn them into plain text files. Non-ASCII characters are filtered out in this step, for the convenience of later analyses. Next, for each manuscript, we segment it into chunks of with length 512 for further Binoculars calculation.

### 4.2. Identification of country/region information

For country/region and language analysis (Fig. 2), at least one country/region and language must be assigned to each manuscript. All platforms do not provide author country/region information directly. bioRxiv and medRxiv provide the corresponding author name and institution information. arXiv provides only a list of author names for each paper.

To simplify and implement our analysis, we made several decisions in this process. First, we assume that corresponding author largely determines the content and writing style of the manuscript. Second, we assume the last author provided by arXiv at most of the time, can be treated as the corresponding author of that paper. bioRxiv and medRxiv do not rely on this assumption as they provide the information directly. for bioRxiv and medRxiv, when the corresponding author is affiliated with more than one institution, we selected the first one as the only institution for our analysis. Third, when the corresponding author’s institution information is available, we use Google Maps API to get the country/region information of the institution and treat this as the country/region of the paper. Lastly, when the corresponding author’s institution information is not available (arXiv), we use nationalize.io’s service to infer the country/region of the corresponding author, as one’s name statistically can be used to infer their ethnicity/nationality (needs citation). Then the official language is determined using the ISO-3166 Country Codes and ISO-639 Language Codes. Since nationalize.io also returns unknown for some names occasionally, we excluded such manuscripts for the country/region/language analysis.

### 4.3. ChatGPT Google Trends

We downloaded worldwide Google Trends data for the keyword “ChatGPT “, both weekly and daily, from https://trends.google.com/trends/. Weekly trends data is downloaded directly from the server. But for the daily trends data, since Google only provides a limited time interval for each query and the results are normalized within the interval from 0 to 100, we downloaded daily data in two-month intervals and forced different queries to overlap. This approach allows the reconstruction of long-interval daily trends data using the earliest month as the base.

### 4.4. Binoculars score

For each manuscript, after segmenting into text chunks with size 512, we used the detector package https://github.com/ahans30/Binoculars provided by (Hans et al., 2024) to calculate the Binoculars score. Specifically, the input text is first tokenized into a sequence of tokens and then fed into two separate LLMs. In our case, we used the Falcon-7b model and the Falcon-7b-instruct model (Almazrouei et al., 2023). This method first calculates the log perplexity (logPPL) of one LLM using the negative average of the logarithm of the next token probability in the input sequence. Next, a log “cross”-perplexity (logX-PPL) for another LLM is calculated using the negative weighted average of the second LLM’s logarithm of the next token probability, with weights provided by the first LLM. The Binoculars score is then defined by dividing the first negative average by the second “cross”-negative average:

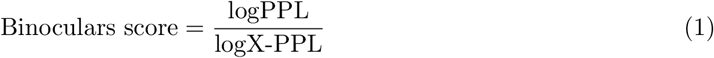

In our analysis, we calculate Binoculars scores for each text chunk in a manuscript. At the manuscript level, we compute the variance, mean, and min values of all Binoculars scores in the manuscript.

### 4.5. Content type classification

To examine the correlation between content types and Binoculars scores, we classified all text pieces into the following 12 content types: phenomenon description, hypothesis formulation, methodology explanation, data presentation, data presentation, logical deduction, result interpretation, literature review, comparative analysis, conclusion summarization, future work suggestions, bibliography, publishing metadata.

This list covers the majority of content found in a typical scientific paper. Subsequently, we employed Meta’s NLI-based Zero Shot Text Classification model (https://huggingface.co/facebook/bart-large-mnli) (Lewis et al., 2019; Yin et al., 2019) to perform zero-shot text classification using the above list of content types. Except for bibliography and publishing metadata, which are not essential for our analysis, the distributions are analyzed.

### 4.6. Regression analysis of language and country/region

We employed Ordinary Least Squares (OLS) regression analysis to investigate the influence of domains and language, and the release of ChatGPT on the min, mean and variance of Binoculars scores of all manuscripts. The models used correspondingly are:

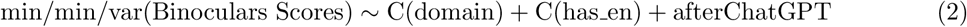

where the domain is from the analysis in Fig. 2A, has en denotes whether the manuscript is from a country/region that has English as one of its official languages (Fig. 4) and afterChatGPT indicates if the manuscript is uploaded after the release of ChatGPT (0: before or 1: after).

**Figure 4.**
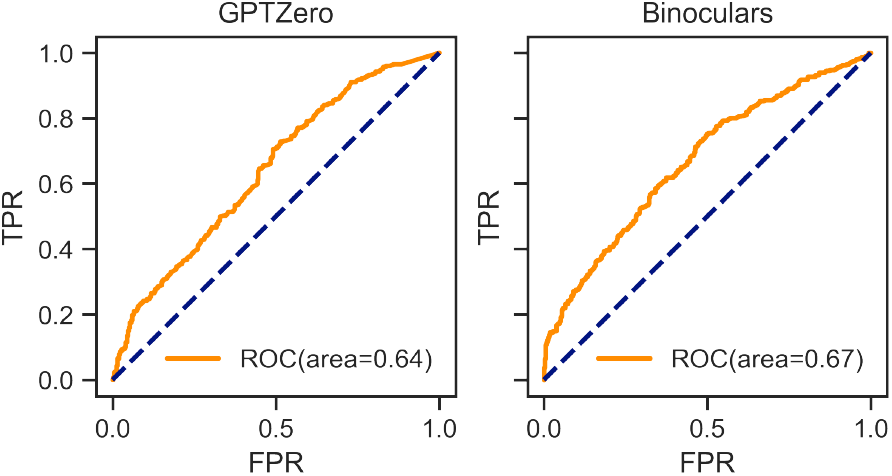
The ROC comparison between GPTZero (left) and Binoculars (right) models.

## A Supplementary Materials

### A.1. Comparison of AI content detectors

We sampled 1,000 manuscripts from those submitted prior to the release of ChatGPT, of which 500 were used as positive samples. For these manuscripts, a maximum of 5 chunk per manuscript were selected to be revised by GPT-3.5, using the following prompt:

You are a helpful assistant. The user will send you a message containing an unoptimized piece of academic writing that is excerpted from a paper. You will revise the piece and improve it. Notice that the piece may be incomplete paragraphs and may have unfnished beginning and ending sentences. You need to respect these parts, do not modify them, and only edit parts that will not influence its original content. Your response should be pluggable to the original paper seamlessly. Your response will be nothing but the modified content. DONOT reply anything else.

This step thus constructed a dataset with ground truth labels of AI revision.

Next, we used Binoculars score detector and an open sourced implementation^3^ of GPTZero to calculate a label for all chunks in each manuscript. Two Logistic regression models are trained based on the label sequence statistics of each manuscript, using the mean, min and variance values. These models predict the presence or absence of AI-revised content in a manuscript. From the dataset of 1,000 manuscripts with ground truth, 80% was used for training and 20% for testing. Both models work with a higher than chance accuracy. The model trained with Binoculars scores achieved an AUC of 0.67, while the one with GPTZero scores achieving a lower 0.64 (Fig. 4).

#### A.1.1 AI usage declaration

We randomly sampled 1000 papers from the manuscripts with lowest 10% average Binoculars scores in the whole dataset. Using GPT and human evaluation, we found no signs of any explicit declarations of AI/LLM/ChatGPT usage in them. Below is the prompt used for initial AI usage declaration:

You are an AI assistant whose role is to analyze academic papers submitted by users in plain text format. Your specific task is to determine whether the paper includes any declarations or statements indicating that it has been edited, revised, or written with the assistance of Artificial Intelligence (AI), Language Models (LLM), or specifically ChatGPT. It is crucial to focus solely on the content generation aspect of writing, excluding any involvement of AI in data preparation, data analysis, or other non-writing related activities. After your analysis, you will respond with a single letter: “Y” for Yes if you find evidence indicating that the paper’s textual content was AI-generated, or “N” for No if there is no indication of AI-generated writing. Your evaluation should be accurate, honing in on explicit acknowledgments of AI’s role in the creation of the paper’s written content.

### A.2. OLS results

To investigate domain/language influence on Binoculars scores, OLS regressions are performed for the mean/min/variance of manuscripts’ Binoculars scores.

**Table 1:** OLS results for mean values of Binoculars scores.

|  | coef | std err | *t* | P>\|*t*\| | [0.025 | 0.975] |
| --- | --- | --- | --- | --- | --- | --- |
| Intercept | 1.0285 | 0 | 2657.55 | 0 | 1.028 | 1.029 |
| C(fields)[T.Computer Science] | -0.0221 | 0 | -54.817 | 0 | -0.023 | -0.021 |
| C(fields)[T.Economics and Finance] | -0.014 | 0.001 | -12.03 | 0 | -0.016 | -0.012 |
| C(fields)[T.Engineering] | -0.0157 | 0.001 | -14.899 | 0 | -0.018 | -0.014 |
| C(fields)[T.Environmental Sciences] | -0.0029 | 0.001 | -2.34 | 0.019 | -0.005 | 0 |
| C(fields)[T.Mathematical Sciences] | -0.0266 | 0.001 | -52.156 | 0 | -0.028 | -0.026 |
| C(fields)[T.Medicine] | -0.0019 | 0 | -5.018 | 0 | -0.003 | -0.001 |
| C(fields)[T.Neurosciences] | 0.003 | 0.001 | 4.598 | 0 | 0.002 | 0.004 |
| C(fields)[T.Physical Sciences] | -0.022 | 0 | -52.336 | 0 | -0.023 | -0.021 |
| C(has_en)[T.1] | 0.0014 | 0 | 6.445 | 0 | 0.001 | 0.002 |
| afterChatGPT[T.True] | -0.0014 | 0 | -6.423 | 0 | -0.002 | -0.001 |

**Table 2:**
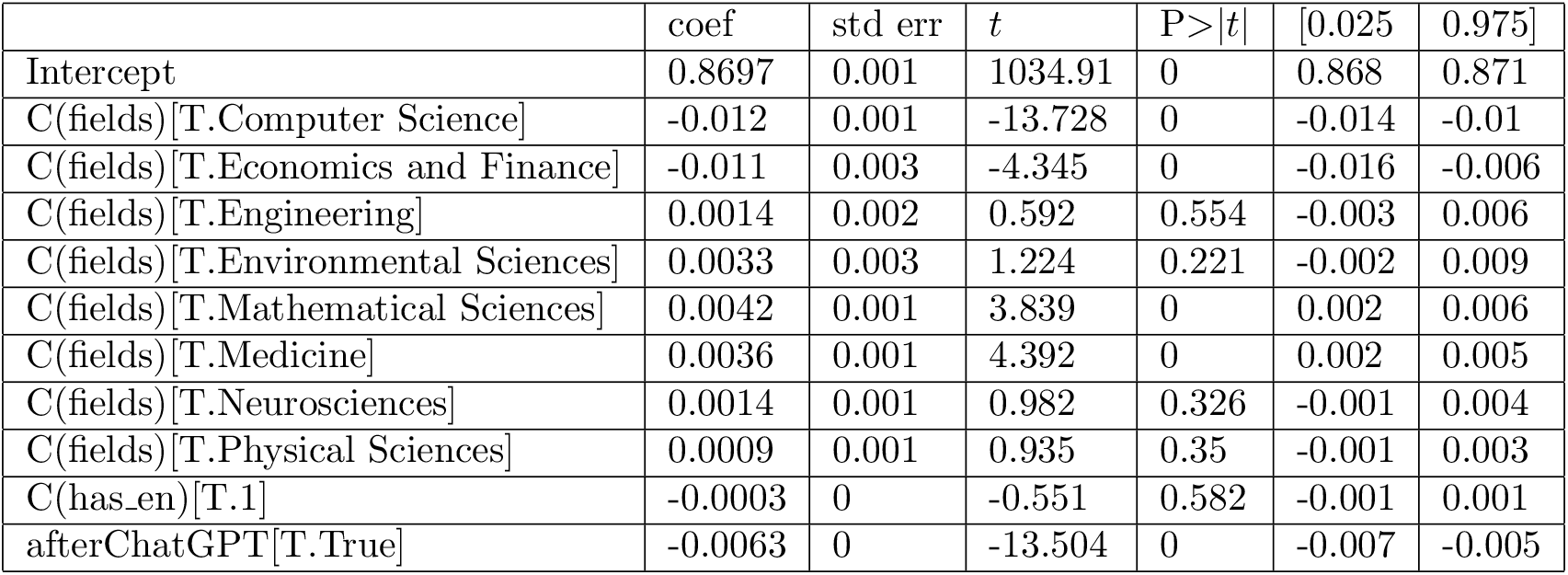
OLS results for min values of Binoculars scores.

**Table 3:**
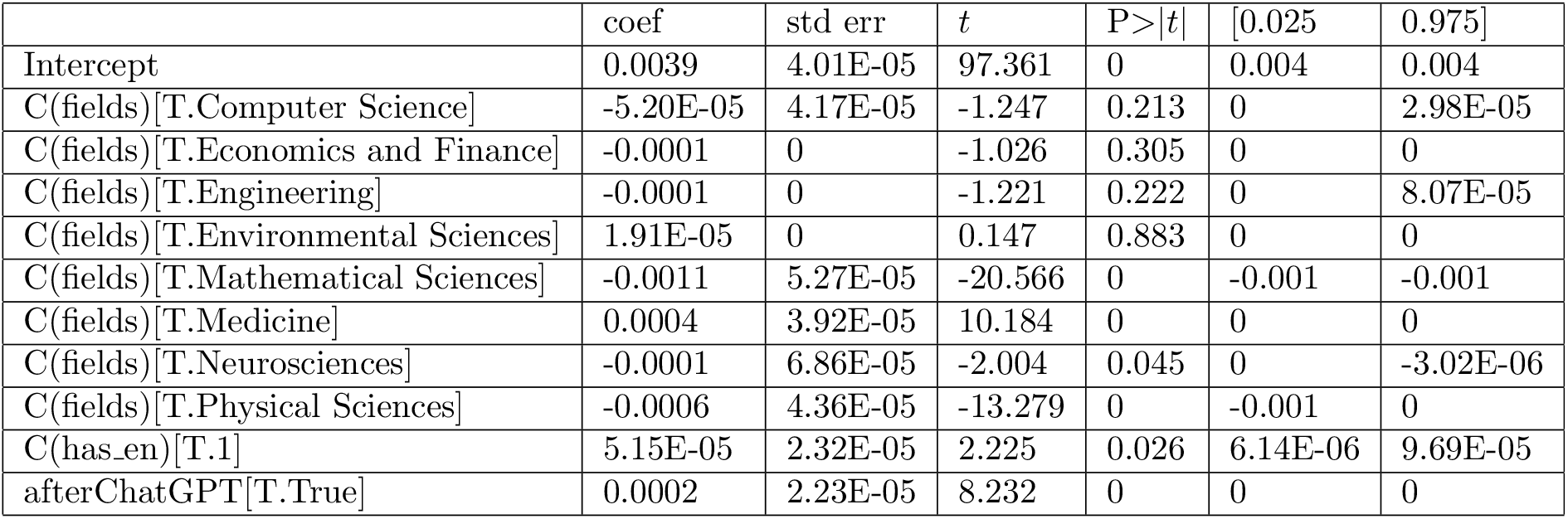
OLS results for Binoculars scores variance.

|  | coef | std err | *t* | P>\|*t*\| | [0.025 | 0.975] |
| --- | --- | --- | --- | --- | --- | --- |
| Intercept | 0.0039 | 4.01E-05 | 97.361 | 0 | 0.004 | 0.004 |
| C(fields)[T.Computer Science] | -5.20E-05 | 4.17E-05 | -1.247 | 0.213 | 0 | 2.98E-05 |
| C(fields)[T.Economics and Finance] | -0.0001 | 0 | -1.026 | 0.305 | 0 | 0 |
| C(fields)[T.Engineering] | -0.0001 | 0 | -1.221 | 0.222 | 0 | 8.07E-05 |
| C(fields)[T.Environmental Sciences] | 1.91E-05 | 0 | 0.147 | 0.883 | 0 | 0 |
| C(fields)[T.Mathematical Sciences] | -0.0011 | 5.27E-05 | -20.566 | 0 | -0.001 | -0.001 |
| C(fields)[T.Medicine] | 0.0004 | 3.92E-05 | 10.184 | 0 | 0 | 0 |
| C(fields)[T.Neurosciences] | -0.0001 | 6.86E-05 | -2.004 | 0.045 | 0 | -3.02E-06 |
| C(fields)[T.Physical Sciences] | -0.0006 | 4.36E-05 | -13.279 | 0 | -0.001 | 0 |
| C(has_en)[T.1] | 5.15E-05 | 2.32E-05 | 2.225 | 0.026 | 6.14E-06 | 9.69E-05 |
| afterChatGPT[T.True] | 0.0002 | 2.23E-05 | 8.232 | 0 | 0 | 0 |

## Footnotes

1 In this study we may use LLM, ChatGPT and AI alternately. As in the use case of text generation, the most advanced AI tools are usually transformer based LLMs, and ChatGPT has been the dominating choice among these LLMs at the time of writing.

2 available from https://gptzero.me

3 https://github.com/BurhanUlTayyab/GPTZero

